# Dendritic cell effector mechanisms and tumor immune microenvironment infiltration define TLR8 modulation and PD-1 blockade

**DOI:** 10.1101/2024.09.03.610636

**Authors:** Daniel A. Ruiz-Torres, Jillian Wise, Brian Yinge Zhao, Joao Paulo Oliveira-Costa, Sara Cavallaro, Peter Sadow, Jacy Fang, Osman Yilmaz, Amar Patel, Christopher Loosbroock, Moshe Sade-Feldman, Daniel L. Faden, Shannon L. Stott

## Abstract

The potent immunostimulatory effects of toll-like receptor 8 (TLR8) agonism in combination with PD-1 blockade have resulted in various preclinical investigations, yet the mechanism of action in humans remains unknown. To decipher the combinatory mode of action of TLR8 agonism and PD-1 blockade, we employed a unique, open-label, phase 1b pre-operative window of opportunity clinical trial (NCT03906526) in head and neck squamous cell carcinoma (HNSCC) patients. Matched pre- and post-treatment tumor biopsies from the same lesion were obtained. We used single-cell RNA sequencing and custom multiplex staining to leverage the unique advantage of same-lesion longitudinal sampling. Patients receiving dual TLR8 agonism and anti-PD-1 blockade exhibited marked upregulation of innate immune effector genes and cytokines, highlighted by increased *CLEC9A+* dendritic cell and *CLEC7A/SYK* expression. This was revealed via comparison with a previous cohort from an anti-PD-1 blockade monotherapy single-cell RNA sequencing study.

Furthermore, in dual therapy patients, post-treatment mature dendritic cells increased in adjacency to CD8^+^ T-cells. Increased tumoral cytotoxic T-lymphocyte densities and expanded CXCL13^+^CD8^+^ T-cell populations were observed in responders, with increased tertiary lymphoid structures (TLSs) across all three patients. This study provides key insights into the mode of action of TLR8 agonism and anti-PD-1 blockade immune targeting in HNSCC patients.

## 1. Introduction

Head and neck cancers are the sixth most common malignancy worldwide [1], and head and neck squamous cell carcinomas (HNSCC) constitute over 90% of these cases. Approximately 50-60% of patients experience loco-regional recurrence within two years of starting therapy [2, 3], underscoring the urgent need for new therapeutic approaches. Immune checkpoint blockade (ICB) activates T cell antitumor activity by targeting immunoregulatory molecules such as the programmed cell death receptor 1 (PD-1). This approach has become the standard of care for recurrent or metastatic HNSCC, as these tumors often have the highest levels of immune cell infiltration for solid malignancies [4]. Despite this, response rates to single-agent anti-PD-1 are low (<18%) [2], and durable complete responses are scarce. Neoadjuvant ICB has shown significant promise in HNSCC, but overall response rates remain modest [5]. Thus, further exploration into mechanisms of neoadjuvant immune targeting in HNSCC is needed.

The activation of Toll-like receptors (TLRs) stimulates innate immune cells, releasing cytokines and enhancing T-cell and dendritic cell (DC) functions. [6]. Dendritic cells are antigen-presenting cells present in the tumor immune microenvironment (TME) of HNSCC [7] and have been widely proposed as promising targets for cancer immunotherapy [8]. TLR agonism and dendritic cell vaccination have shown increased interferon gene expression and prolonged survival in a randomized phase II clinical trial in malignant glioma [9]. Recent work in HNSCC mouse models has shown the ability of DCs to overcome resistance to ICB by enhancing T-cell infiltration after conventional dendritic cell vaccination [10], underscoring the potential role of DCs in ICB success.

Within the TLR family, TLR8 recognizes single-stranded RNA molecules and leads to the induction of a type I interferon response, which results in the production and secretion of proinflammatory cytokines and chemokines (e.g., CXCL9, CXCL10). Recently, motolimod (VTX-2337), a selective synthetic TLR8 agonist, has been tested as adjuvant therapy in combination with chemotherapy in HNSCC patients, showing promising results, particularly in human papillomavirus (HPV) positive patients [11]. Motolimod’s properties may rely on understudied distinct TMEs, and to the best of our knowledge, its effects on the transcriptional phenotypes of the malignant cells and adjacent TME have never been studied in humans.

Prior preclinical work on TLR stimulation has shown that CD11c-expressing dendritic cells are activated to enter an organized non-lymphoid tissue aggregate known as a tertiary lymphoid structure (TLS) [12]. TLR agonism also contributes to the formation of TLSs and the upregulation of CXCL13 in myofibroblasts and lung tumors [13,14]. Importantly, TLSs have also been revealed as a surrogate marker of a robust immune response and are associated with improved responses to ICB, yet CXCL13 serum levels are a poor prognostic marker in certain lung cancers [15,16]. Thus, the composition and spatial characterization of TLSs could provide a better understanding of the impact of TLR8 agonism and PD-1 blockade in HNSCC patients.

Advancements in single-cell genomics have provided an avenue for capturing the heterogeneity of epithelial tumors and their microenvironment. Single-cell RNA sequencing (scRNAseq) studies have revealed that DCs decrease in the HNSCC tumor microenvironment post-ICB therapy [17].

Multiplex-immunofluorescence (mIF) imaging has been proven to be an effective tool for biomarker discovery and to predict responses to ICB in different solid malignancies, including HNSCC [18].

Given myeloid cells’ importance in driving antitumor activity, the decrease of DCs post- ICB therapy, and the association of TLSs with response to ICB, we set out to determine the effect of TLR8 agonist (motolimod), in combination with anti-PD-1 blockade (nivolumab) on the tumor ecosystem in HNSCC patients. As part of a phase 1b clinical trial (NCT03906526), we used single-cell sequencing alongside multiplex immunofluorescence (mIF) imaging tools to analyze pre- and post-treatment tumor samples taken from the same lesion to investigate this dual mode of action. Our study design allowed us to profile TME changes across treatments, identifying dynamic transcriptional signatures in the DC population that were unique to dual therapy and were not present in a patient cohort treated with anti-PD-1 blockade monotherapy alone.

## Materials and Methods

### Tissue collection, processing, and response assessment

Patients with previously untreated, resectable HNSCC were recruited as part of a clinical trial https://clinicaltrials.gov/ct2/show/NCT03906526. Tissue genomic profiling was approved by the Dana-Farber/Harvard Cancer Center Institutional Review Board (DF/HCC Protocol # 21-312). Briefly, fresh tumor tissue biopsies were collected before two doses of anti-PD-1 blockade (nivolumab) and four doses of subcutaneous (SC) toll-like receptor 8 (TLR8) agonist (motolimod). The tumor was then surgically resected (days 23, 30, and 28 for patients 1, 2, and 3, respectively) (**Figure 1A**). Response assessment was evaluated by a head and neck neuroradiologist according to RECISTv1.1 criteria (**Supplementary Table 1**). Patients are designated by their response as responders: Pt1_R and Pt2_R, and non-responder: Pt3_NR.

**Figure 1.**
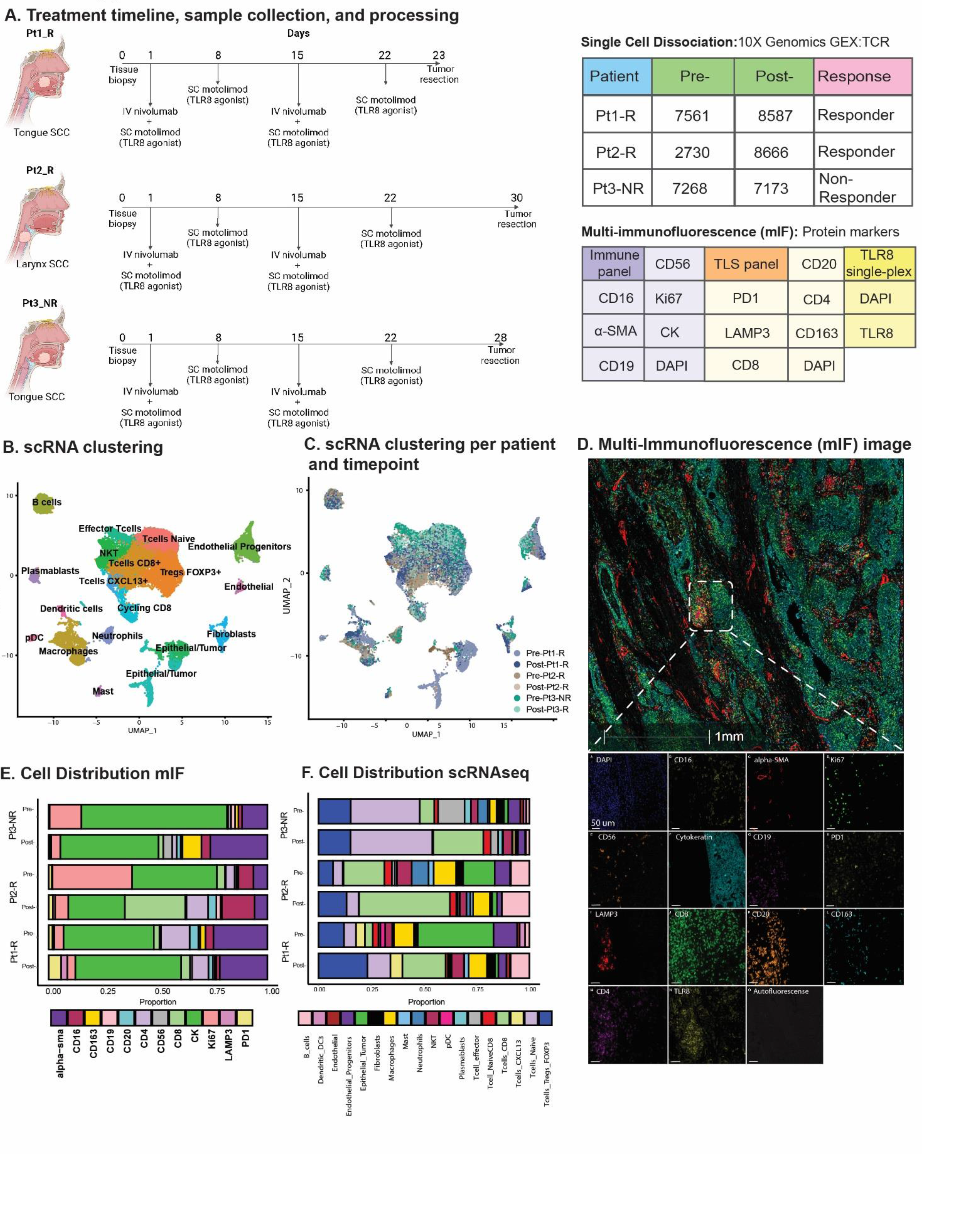
Overview of TLR8 agonism and anti-PD-1 blockade workflow and cell types identified. (A) Schematic representation of treatment and biopsy collection created with BioRender.com. The tables show the number of cells recovered after scRNAseq processing, patient biopsy site, patient response, and antibodies used for multi-immunofluorescence (mIF) imaging. **(B)** UMAP shows the clustering of all 41,985 cells collected in this study. **(C)** UMAP shows the patient contribution to the twenty-two clusters identified in B. **(D)** Representative image showing the mIF image of post-treatment biopsy of Pt2_R. **(E-F)** Cell proportions found by mIF imaging **(E)** and by scRNAseq **(F)** for all patients pre- and post-treatment.

### Isolation of tumor and immune cell infiltrates for scRNAseq

All tumors were dissociated using the Human Tumor Dissociation kit (Miltenyi Biotec; Cat#130-095-929) within 30 minutes after surgery (**Supplementary Methods**) and were prepared as previously described [19]. Briefly, minced fragments were incubated with DMEM, enzyme H, enzyme R, and enzyme A and vortexed for 15 minutes at 400 rpm, 37°C. Tissue was filtered (50µm filter, Sysmex; Cat# 04-004-2327), washed, minced with a 1mL syringe plunger, and re-washed. Dissociated cells were filtered and spun again (30µm filter, Sysmex; Cat# 04-004-2326) at 1500 rpm at 4°C for 5 minutes. Next, incubation with ACK buffer (Gibco, Cat#A1049201) was performed, followed by removal of the ACK buffer. In all cases, the viability of dissociated single-cell suspensions was increased to >=90% by using a modified protocol of the EasySep Dead Cell Removal (Annexin V) kit (STEMCELL; Cat#17899). Gene expression (GEX) and TCR libraries were generated using the 10x Genomics Chromium NextGEM Single Cell V(D)J Reagents kits v1.1 (PN-1000165; PN-1000020; PN-1000005; PN- 1000120) according to the manufacturer instructions.

#### FFPE tissue specimens

Serially cut 4-µm thick sections were obtained from Formalin Fixed Paraffin Embedded (FFPE) tumor biopsies. Human tonsils, and discarded HNSCC FFPE tissue blocks were used for conventional immunohistochemistry (IHC), multiplex validation, and assay optimization. Before staining, all tissue slides were deparaffinized and rehydrated by serial passage through graded concentrations of ethanol. Briefly: slides were submerged in xylene (Millipore XX0060-A) during 10 min (x3), quick wash in deionized (DI) water, 100% ethanol x 10 min, 95% ethanol x 10 min, 70% ethanol x 5 min, quick wash in DI water, 10% neutral buffered formalin (Thermo Scientific Cat#5725) x 15 min, followed by 5 min of running water before antigen retrieval in Tris EDTA buffer pH 9.0 (Vector Laboratories H-3301-250) or Citrate-based pH 6.0 (Vector Laboratories H-330-250) buffer (exclusively for anti-CD20 antibody).

### Multiplex Immunofluorescence (mIF) staining

After immunohistochemistry validation and spectral library creation (**Supplementary Methods**), we utilized a commercially available manual mIF staining kit (Opal 7-plex, Akoya, Cat# NEL811001KT) and applied an optimized protocol to all mIF staining (**Supplementary Table S2**). Serial manual staining was performed as suggested by the manufacturer (https://www.akoyabio.com/wp-content/uploads/2020/04/Opal-Reagents_Brochure_Opal-Assay-Development-Guide.pdf). Two panels were optimized for staining, Immune Panel (Panel A) and TLS panel (Panel B). Two serial sections were used, each stained with one panel. An additional slide was later used to identify TLR8 expression in tissue. This slide was stained with TLR8 and DAPI, scanned, and subsequently added to the analysis (**Supplementary Table 2**). The whole tissue section was scanned using our multiplex immunofluorescence (mIF) imaging platform (Vectra 3, Perkin Elmer). Phenochart was used to select all the regions of interest (ROI) for higher-resolution scanning (20x). Spectral unmixing was done using InForm 2.5.1, and unmixed component mIF images (QPTiff format) were then uploaded to the analysis platform (Halo software; Indica Labs).

### Downstream mIF Image Processing

Serially cut stains were registered, and a synchronous navigation tool was used to check fusing quality (**Supplementary Figure S1**). Tumor-stroma classification with Hematoxylin Eosin (H&E) was performed by a head and neck pathologist (PS), and then mIF slides were classified by a random forest classifier trained with pathologists’ annotations. Single-cell phenotyping was done using Indica Labs – HighPlex FL v4.1.3 module (**Supplementary Table S3**). The same thresholds were applied to all the slides, and spatial analysis was performed based on average values. A paired t-test was used to compare pre- and post-treatment values. The peritumoral region was defined as the area extending 100 μm outside the tumor margin, as previously described [20].

### Tertiary Lymphoid Structures quantification

Tertiary Lymphoid Structures were defined as aggregates of lymphoid cells with histologic features resembling follicles in lymphoid tissue. A head and neck pathologist (PS) identified these structures on serially cut H&E slides. Additionally, TLSs were identified on serially cut slides stained with mIF panels using random forest classifiers as described above. At single-cell mIF phenotyping, we included TLSs with more than 50 B cells (CD20+) and more than 5 dendritic cells (LAMP3+). The number of TLSs was normalized by the area analyzed (TLS/mm^2^).

#### Processing of GEX and TCR scRNAseq

Raw sequencing reads were aligned, and samples aggregated without normalization (Cell Ranger v6.1.1). Cells with fewer than 150 genes or 15% of unique molecular identifiers (UMIs) mapped to mitochondrial genes were filtered (Seurat (v3)).

Potential doublets were filtered using DoubletFinder (chris-mcginnis-ucsf/DoubletFinder). Filtered TCR alpha and beta, contigs were then integrated by barcode with scRepertoire (ncborcherding/scRepertoire) based on the variable, diversity, and joining (VDJ) genes and CDR3 nucleotide sequence. 10X genomics feature matrices from Luoma et.al [17], representing anti-PD-1 monotherapy, were accessed from Gene Expression Omnibus Accession (GSE200996) and processed as described above (**Supplementary Methods**).

### Clustering and annotation of tumor and tumor-infiltrating immune cells

Principal component analysis used 2000 highly variable genes from scaled and log normalized expression matrices. Clustering utilized the first 40 principal components with a Louvain algorithm. The annotation of the resulting 22 clusters was based on differential gene expression (log fold threshold: 0.25). T cells and myeloid cell clusters gene expression and TCR clonotypes were extracted for further investigation (**Supplementary Methods**). To complete this work, we also used a published dataset from Luoma et al. [17]. Datasets were integrated by ranking 2000 variable features. 40 principal components were used to identify clusters in the integrated dataset. The annotation was performed based on differential gene expression analysis in combination with the integration of previously labeled clusters from Luoma et al. [17].

### Copy number calling and phylogenetic tree construction

InferCNV (InferCNV of the Trinity CTAT project: https://github.com/broadinstitute/InferCNV) was used to predict copy number alterations with CD45+ cell clusters as references. Phylogenetic trees of tumor cells were then constructed using Uphyloplot2 (harbourlab/uphyloplot2; **Supplementary Figure S2-4, Supplementary Table S4**).

### Pre- and post-treatment dendritic cell differential gene expression analysis

The gene expressions of extracted myeloid cell clusters from our cohort and the Luoma anti-PD-1 monotherapy cohort [17] were separately log-normalized and scaled, and 2000 variable features were defined. After integrating both myeloid cell matrices, the first 8 principal components were used for clustering. Differential gene expression analysis alongside previous labeling from Luoma et al. [17] provided cluster annotations. Differential gene expression for pre- and post- treatment subset data was performed on cell clusters identified as non-plasmacytoid dendritic cells (LAMP3+ DC, CLEC9A+ XRC1+ DC, LAMP3+ CCL17+ DC, and CD1C+ DC) using a hurdle model (MAST) [21]

(fold change cutoff: 0.01). Gene set enrichment was performed against MSigDB GO Biological Ontology (c5.go.bp.v2022.1.Hs.symbols.gmt) using the fgsea library in R (v4.2.1).

## Data Availability

Processed scRNAseq data is available in the Gene Expression Omnibus (GEO) as h5 files (GSE273138). The scripts used in this study are available at https://github.com/jfwise/HSNCC_Single_Cell_TLR8.

## Results

### Single-cell RNA-sequencing and spatial analysis of HNSCC patients treated with TLR8 + anti-PD-1 combinatorial therapy

To define the effect of combined TLR8 agonism and anti-PD-1 blockade in humans, HNSCC patients with previously untreated, clinically diagnosed, resectable tumors were recruited for an open-label, phase 1b pre-operative window of opportunity biomarker trial (NCT03906526). From combination therapy arm 4, three patients receiving nivolumab (anti-PD-1 blockade) IV for 2 doses on days 1 and 15, and motolimod (VTX-2337/TLR8 agonist) SC injection weekly for four doses on days 1, 8, 15, and 22, had same lesion tumor biopsies, collected pre- and post-treatment. To determine the immunomodulatory mode of action, we employed scRNAseq with T-cell clonotype identification to leverage high-resolution insight into immune cell population dynamics. However, given the loss of tissue architectural elements, we also applied a customized staining protocol (mIF) for spatial organization details of critical immune cell subsets for three patients with the same lesion-matched pre- and post-treatment biopsies (**Figure 1A**). From the scRNAseq sample collection, 41,985 cells were analyzed after applying initial quality controls and doublet removal (**Supplementary Methods**). We partitioned the cells into 22 clusters by their expression states and annotated them using differential gene expression of known markers (**Figure 1B-C; Supplementary Figure S2, Supplementary Table-S5**). To further classify epithelial tumor cells, we identified copy number variations using InferCNV and phylogenetic tree reconstruction of subclones, highlighting the diverse heterogeneity of HNSCC (**Supplementary Figure S2-4, Supplementary Table S4**).

### scRNAseq and mIF identify post-treatment changes in TLR8 expression and TME composition

Cell proportion estimates were calculated by scRNAseq and a serial multi-immunofluorescence (mIF) imaging technique (**Figure 1D-E**; **Supplementary Table S6**). For our patient cohort, two patients responded to therapy (Pt1_R and Pt2_R), while one was identified as a non-responder (Pt3_NR). For all patients, scRNAseq analysis showed a decrease in epithelial tumor proportions post-treatment. However, using mIF, a decrease in epithelial tumor proportions was observed for Pt2_R and Pt3_NR, while an increase was found in Pt1_R (**Figure 1E**). Similar to previous investigations of anti-PD-1 blockade, the total percentage of T-cells increased post-therapy via scRNAseq (**Figure 1E**). Since innate immune cells are the target of TLR8 modulation, we were surprised to see a decrease in the total myeloid population post-treatment, as shown by scRNAseq (**Supplementary Figure S5 A**). However, mIF revealed an increase in dendritic cells (LAMP3+). Importantly, while scRNAseq could not detect TLR8 transcripts, TLR8 was detected at the protein level (**Figure 1D**). We discerned a tendency for higher average TLR8+ cell signal intensity (13, 82 and, 67% increase for Pt1-R, Pt2-R, and Pt3-NR respectively) (**Supplementary Figure S5 B, C**), and a higher percentage of cells positive for TLR8 in post-treatment biopsies (Pt1-R pre: 38.3%, post: 74.41%; Pt2-R pre:54%, post: 57%; Pt3-NR pre: 33%, post: 72%);(**Supplementary Figure S5 D**), mainly co-expressed in CK+ cells (**Supplementary Figure S5 E**). These results highlight the effect of combined TLR8 agonism and anti-PD-1 blockade on the TME in HNSCC.

### Spatial analysis of macrophages and T cells across therapy

Macrophages are essential agents in antitumor immunity but can also have immunosuppressive features. Hence, we examined M2 macrophage spatial behavior via the CD163 marker [22].

Interestingly, the non-responder (Pt3_NR) showed a higher density of CD163+Ki67+ macrophages post-treatment, alongside increased non-proliferating macrophages in the peritumoral space (**Supplementary Figure S6 A-C**). However, we observed no difference in CD163 expression in M2 macrophages via scRNAseq based on response to therapy (**Supplementary Figure S6 D-E)**.

T cells, key in antitumor immunity, activate after antigen recognition by antigen-presenting cells (APCs). Anti-PD-1 blockade is designed to help release restraints on T cell activation. Post-treatment, CD4+ helper cells were in closer proximity to CD8+ cytotoxic T cells in all biopsies, while CD19+ B lymphocytes were closer exclusively in responders (**Supplementary Figure S7**). These results suggest that effective combinatorial therapy may be evidenced by T cell activation supported by helper T cells, and that T-B cell interactions may have an important role in their response. The spatial distribution between T cells and M2 macrophages was inconsistent across all patients (**Supplementary Figure S7**). An increase in peri- and intra-tumoral CD8+ cytotoxic T cells was observed among all patients in post-treatment biopsies, with a higher proportion in responders (**Figure 2A: Supplementary Figure S8 A**). Increased T-cell density in the peritumoral area may result in increased exposure to neoantigens through T-cell receptor (TCR) engagement and subsequent T-cell clonal expansion.

**Figure 2.**
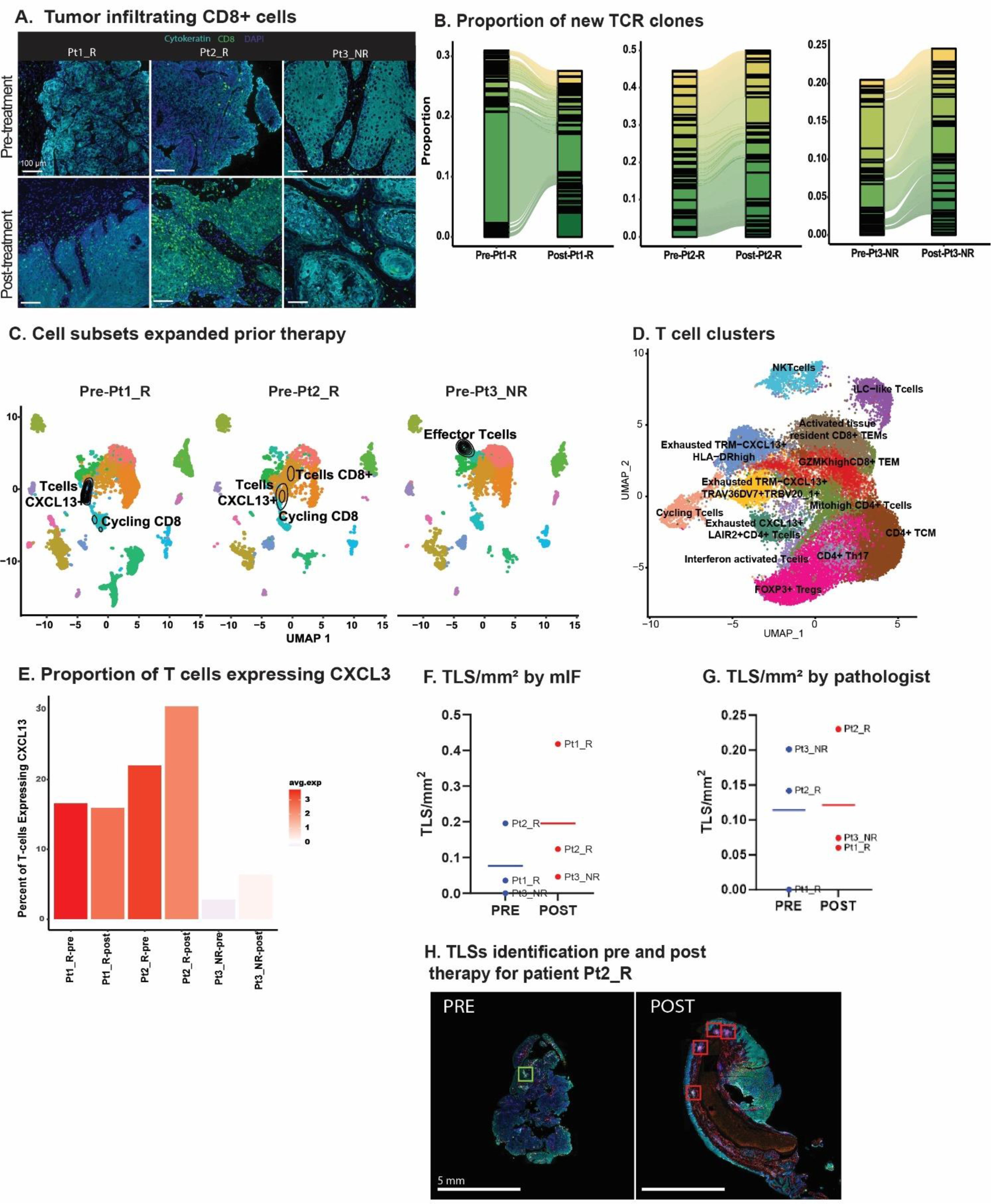
TLR8 agonism and anti-PD-1 blockade increase TCR clones and Tertiary Lymphoid Structures (A) Representative image showing tumor infiltrating CD8 T cells (green) before and after anti-PD-1 blockade and TLR8 agonism for each patient. **(B)** An alluvial plot depicts the proportion of new TCR clones post-treatment per patient; only the top thirty clones are represented per patient. **(C)** A UMAP of each patient’s pre-treatment biopsy with the density of cells with a TCR expansion >10 depicts which cell subsets expanded prior to TLR8 agonism and anti-PD-1 blockade. **(D)** UMAP displays the 13 T cell clusters identified after isolation and sub-setting of prior recognized T and NKT clusters. **(E)** A bar plot with varied colors reveals the average gene expression and percentage of *CXCL13-*expressing cells across T cells. **(F)** Number of TLS/mm^2^ identified by mIF. **(G)** Number of TLS/mm^2^ identified by the pathologist. **(H)** Representative image of tertiary lymphoid structures in pre- and post-treatment biopsies from Pt2_R.

### CD8 T-cell clones expand in response to treatment

Given the assumed activation of APCs by TLR8-agonism and increased T-cell density in areas enriched with B cells, we investigated TCR expansion through single-cell TCR-VDJ sequencing. Examination of the frequency and cluster distribution of TCR clonotypes, defined as those TCR sequences identical across more than one cell, revealed an increase in TCR clonotypes post-treatment (**Supplementary Figure S8 B**). Moreover, there was a substantial increase in new TCR clones in post-treatment biopsies (**Figure 2B**). We observed that the majority of expanded TCR clones resided in clusters previously identified as CD8+ T-cells (**Supplementary Figure S8 C, D**). Interestingly, in the pre-treatment biopsy of the non-responder, there was an expansion in ILC1-like T-cells (*GZMH+*, *NKG7+*, *GNLY+)*, while the majority of TCR expansion in responders occurred within cycling CD8+CXCL13+ T cells, a chemotactic for B-cells (**Figure 2C**).

To further refine TCR clonotypes and investigate the contribution of CXCL13+ expression in TCR expansion, we partitioned the seven clusters identified as T cells. We identified 13 clusters (**Figure 2D**; **Supplementary Table S7**). Two clusters had little CD4 or CD8 expression, representing an NKT cell cluster and the previously described ILC1-like T cells (**Supplementary Figure S8 E**). Four clusters (Cycling:Exhausted TRM-CXCL13+HLA-DRhigh: Exhausted TRM- CXCL13+TRAV36DV7+TRBV20_1+: Exhausted CXCL13+LAIR2+CD4+ T cells) had a differential expression of *CXCL13* including a CD4 positive cluster. Intriguingly, all four clusters differentially expressed exhaustion markers *HAVCR2* and *ENTPD1* (**Supplementary Figure S8 E**). We further investigated *CXCL13* expression across all patients and saw that both, the average expression and the percentage of cells expressing *CXCL13* increased in all patients post-treatment, yet the non-responding patient had overall lower levels of *CXCL13* (**Figure 2E**).

### Tertiary Lymphoid Structures increase after treatment

Next, we evaluated TLSs presence and composition in our patient’s pre- and post-treatment biopsies. This effort was initiated after observing CD4+ and CD8+ T cells closer to CD19+ B cells post-therapy. Further, the role of *CXCL13* in TLS formation has been linked to improved response to ICB [23, 24]. TLSs were defined using criteria established by others [15], with the additional inclusion of lymphoid follicles with germinal centers (>50 CD20+ B cells, and >5 LAMP3+ dendritic cells). With these criteria in place, random forest classifiers were used for mIF TLSs detection (**Supplementary Figure S9 A, B**). Normalized by tissue size, post-treatment biopsies showed a trend of increased TLS/mm2 via mIF (pre: 0.09 TLS/mm2; post: 0.27 TLS/mm2; **Figure 2F**); this trend was confirmed on H&E slides from patients Pt2_R, and Pt3_NR, but not Pt1_R as reviewed by a pathologist (PS) (**Figure 2** **G, H; Supplementary Figure S9 C).** Interestingly, on average, for all pre-treatment biopsies, helper T cells (CD4+) were the primary cell type in TLSs (41%), followed by CD20+ B cells (32%; **Supplementary Table S8 D**). For all post-treatment biopsies, TLSs were largely comprised of cytotoxic T cells (CD8+) (47%), and PD-1 positivity did not increase, although dendritic cells (LAMP3+) did (**Supplementary Figure S9 D; Supplementary Table S8**). No difference was found in their distance to the tumor from pre- to post-treatment biopsies (**Supplementary Figure S9 E**).

Literature on dendritic cell infiltration into TLSs using pre-clinical models, particularly with TLR stimulation, has been contradictory [25], although it is hypothesized that their antigen presentation capabilities enhance T-cell mediated ICB response [26].

### Unique dendritic cell activation in the context of combinatorial therapy

We then aimed to define dendritic cell activation by TLR8 agonism plus PD-1 blockade compared to PD-1 blockade alone. Although we did not have access to samples from this arm of the clinical trial, we utilized a publicly available scRNAseq dataset of three pre- and post-anti-PD-1 treated HNSCC patients as controls (**Figure 3A**) [17]. After integrating the two datasets, we defined 11 distinctive myeloid clusters (**Figure 3B**; **Supplementary Tables S9-10),** including *CD14*+ and *FCGR3A+* monocytes. Two tissue-associated macrophage populations were defined by *ITGAX* expression, cytokine IL-8 producing *CXCL8,* and EMT-related *SPP1* (**Supplementary** Figure 10A). Inflammatory CCL18+ M2-like macrophage subset was defined by *CD206*, *CD163*, and *CD209*. One cluster exhibited *CD3* expression, suggesting a myeloid/T-cell interaction or doublet with failed identification by doublet filtering tools. Thus, we did not investigate this cluster further. The remaining clusters represented five dendritic cell (DC) populations, including plasmocytoid DCs, conventional DCs (*CD1C)*, two mature LAMP3+ DC subsets containing markers for Th2 chemoattractant, *CCL17*, and cell migration factor *FSNC1.* Prior investigations revealed decreased CD1C+ conventional and LAMP3+ DCs with ICB therapy. Importantly, dual-immune targeting of TLR8 and PD-1 revealed an increase in conventional dendritic cells as well as dendritic cells expressing *CLEC9A+* and *XCR1* (**Figure 3C**).

**Figure 3.**
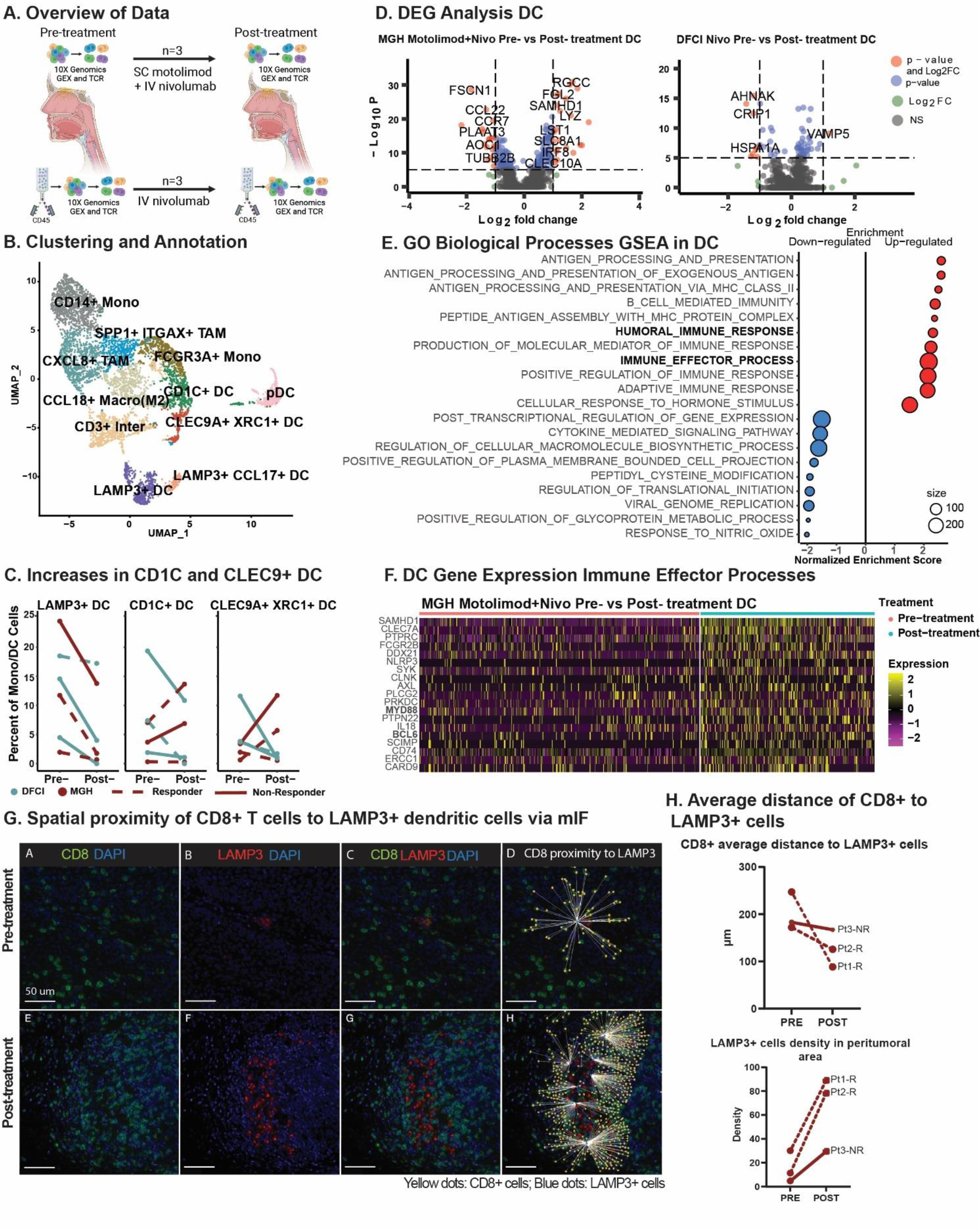
Dendritic cells effector activation in combinatorial treatment compared to monotherapy and spatial dendritic cell proximity to T cells. (A) Schematic representation of data processing and comparison between our (top) and Luoma et al. (bottom) single-cell RNA datasets, created with BioRender.com. (B) A UMAP displays the clustering and annotation of selected myeloid cells after merging the two myeloid cells from the two cohorts. (C) Percent of mature LAMP3+, conventional CD1C+, and CLEC9+XRC1+ dendritic cells. (D) A volcano plot displays the differentially expressed genes of non-plasmacytoid dendritic cells pre- vs. post-treatment in the MGH and Luoma et al. cohorts. (E) Gene Set Enrichment Analysis from the gene ontology biological processes of differentially expressed genes in non-plasmacytoid dendritic cells from the MGH cohort in pre- and post-treatment cells was performed, and a dot-plot displays the enrichment score with color representing the direction of expression for the gene sets, and size represents the number of genes involved in the analysis. (F) Heatmap display of the relative expression of selected genes from the humoral immune response gene sets for non-plasmacytoid dendritic cells pre- and post-treatment for our cohort. (G) Representative image of Pt_2_R showing the spatial behavior of CD8+ T cells (green) and LAMP3+ DC (red) pre-and post-treatment. (H) Average distance of CD8+ T cells to LAMP3+ dendritic cells (upper plot) and LAMP3+ dendritic cell density in the peritumoral area (lower plot).

To further explore the effects of TLR8 agonism and PD-1 blockade in DCs, we performed differential gene expression analysis of non-plasmacytoid DC populations. With dual therapy, we revealed multiple significant gene-level changes in DCs but few significant changes with anti-PD-1 monotherapy (**Figure 3D**). Gene-set enrichment revealed that up-regulated genes are involved in humoral and immune effector responses, including MHC Class II (**Figure 3E; Supplementary Figure S10**), signaling proteins *MYD88*, *SCIMP,* and interferon associated signaling *AXL* and *SAMHD1*. Positive cytokine regulation was also seen with the up-regulation of *CLEC7A* and *SYK*. Moreover, post-treatment, we identified down-regulated genes that have a role in cytokine signaling, including significant decreases in *CCL17* and *CCL22,* suggesting a potential decrease in Th17 and Treg recruitment. A decrease in *CCR7* suggests decreased migration to lymph nodes (**Figure 3F**; **Supplementary Figure S10**). We investigated whether changes in cytokine signaling and MHC Class II activation resulted in shifts in the spatial dynamics of myeloid cells within the TME using mIF. Our analysis revealed a closer proximity between proliferative DCs (LAMP3+Ki67+) and natural killer cells (CD56+CD16-) in the post-treatment biopsies (pre: 251.58 µm; post: 137.37 µm; **Supplementary Figure S7**). Additionally, cytotoxic T cells (CD8+; pre: 200.62 µm: post: 127.03 µm) were found to be closer to DCs (LAMP3+) in all post-treatment biopsies (**Figure 3G**).

Furthermore, a high density of DCs (LAMP3+) was found in the peritumoral area in post-treatment biopsies (paired t-test; p-value 0.0601, **Figure 3H**), indicating a trend toward tumor-specific dendritic cell activation after dual therapy.

## Discussion

In this distinctive window of opportunity phase 1b clinical trial setting, we characterized the effect of TLR8 agonism and anti-PD-1 blockade in the TME of three patients with treatment-naive HNSCC. To accomplish this, we performed single-cell transcriptomic profiling together with high-plex tissue imaging, enabling us to obtain a high-resolution understanding of immune cell responses while preserving the spatial and architectural elements of the TME. Of the patients analyzed, two responded to treatment, while one did not respond. The two responding patients exhibited clonal expansion of *CXCL13*+ exhausted tissue-resident memory CD8+ T cells, suggesting successful reinvigoration of CD8+ T cells and antigen stimulation, as shown by scRNAseq analysis [27]. Consequently, a higher density of tertiary lymphoid structures was found in post-treatment biopsies using machine learning and mIF. The ability to identify and characterize an increased number of TLSs highlights the potential for novel digital pathology methods, such as mIF, to assist with the rapid and reliable identification of these complex structures. These findings are aligned with prior conclusions that correlate *CXCL13* and the presence of TLSs with improved anti-PD-1 response in several solid malignancies [23]. Given our finding of increased LAMP3+ dendritic cells, we used scRNAseq to compare a prior study of anti-PD-1 alone [17] regimens with our combinatorial TLR8 agonism and revealed an exclusive increase in the percentage of conventional DCs (CD1C+) and CLEC9A+XRC1+DCs, which excel at cross-presentation and cytotoxic T lymphocyte responses [28]. Moreover, we observed a humoral and innate effector immune response uniquely in the DC populations exposed to TLR8 agonism and anti-PD-1 blockade (scRNAseq). Lastly, to further investigate the dendritic cell findings from scRNAseq we identified a closer localization of LAMP3+ DC to CD8+ T cells after combinatorial treatment (mIF). Our findings highlight the potential role of activated and mature DCs interaction with CD8+ T cells as critical surrogates of TLR8 agonism and anti-PD-1blockade in a human clinical trial setting, which should be further investigated with increased patient numbers.

TLR8 expression is thought to be enriched in neutrophils, monocytes, dendritic cells, and macrophages. While we were unable to capture *TLR8* RNA expression, and our protein expression was mainly associated with epithelial cells, we revealed major shifts in innate cell populations in the three patients post-treatment. LAMP3+ cells marking mature dendritic cell populations were found closer in distance to CD163+ cells in the non-responder as opposed to CD8+ T-cells in responders (mIF). Additionally, we did not see an increase in tumor-infiltrating DCs by scRNAseq, however, we were able to identify a distinct DC activation following TLR8 agonism and anti-PD-1 blockade.

After comparing our results with an available scRNAseq dataset of HNSCC patients of similar size (n=3) treated with anti-PD-1 monotherapy [17], we saw an increase in dendritic cells associated with T-cell responses and antigen presentation (CD1C+ and CLEC9A+ XCR1+ DCs) in two of three patients. This increase imparted a distinct upregulation of immune effector genes in non- plasmacytoid DCs (scRNAseq) and a clear localization of LAMP3+ DC to CD8+ T cells after treatment (mIF). Prior reports have shown functional localization of T cells around mature DC as a prognostic factor [29]. Indeed, a high peritumoral density of mature DCs has been associated with significantly longer survival in solid malignancies, including esophageal squamous cell carcinoma [30], highlighting DCs as a key component of antitumor activity.

Despite using scRNAseq and cell-to-cell interactions through an image registration machine-learning algorithm, our study has limitations. Our findings were determined from a small subset of one arm from our clinical trial (n=3), and scRNAseq dendritic cell findings required the additional use of publicly available data (n=3) to analyze our cohort. Moreover, despite being a target of TLR8 agonism, we did not see an increase in TLR8 signaling molecules (except *MYD88* in dendritic cell populations) in post-treatment biopsies, possibly because of tumor resection timing (scRNAseq).

Considering motolimod half-life of 6 hours [31], it is possible that TLR8 agonism was not enough to generate a sustained myeloid cell intra-tumoral activity, that we were unable to capture the initial signaling pathway, or that the main location for APC activation took place in the lymph node. While analyzing pre-post drug tissue biopsies is a powerful approach, immediate post-treatment local effects remain poorly assessed with such an approach. Negative feedback after stimulation might explain the absence of TLR8 RNA-based gene expression despite the upregulation of downstream targets such as *MYD88* and *BLC6* [32]. Future investigations into early (<24h) and acute (<2 weeks) post-treatment TME states will be of interest. Moreover, due to our sample size, we were unable to decipher whether TLSs formation increased due to TLR8 agonism or simply by anti-PD-1 blockade alone. Lastly, we could not perform statistical correlates with response due to the small cohort size. Despite the limited sample size afforded by this unique clinical trial, we were able to perform a deep and detailed analysis per sample at the single-cell level (RNA and protein), which identified unique dendritic cell molecular and spatial changes after combined therapy with TLR8 agonism and anti-PD- 1 blockade and indicate that further investigation with a larger sample size is warranted.

## Author Contributions

S.L.S., D.L. F., and M.SF. conceptualized the investigation and provided supervision, and D.L.F. led the funding acquisition. Formal analysis, including visualization, was performed by D.A.RT., J.F.W, B.Y.Z, and P.S, J.F processed the single-cell RNA sequencing samples. S.C., J.P.O.C., and P.S. provided methodology and software support. C.L. and A.P. assisted with study design and sample acquisition. O.Y. provided essential resources. D.A.R.T. and J.F.W. wrote the original draft, and all authors performed critical reviews and revisions.

