## Supplementary Material for "Dendritic cell effector mechanisms and tumor immune microenvironment infiltration define TLR8 modulation and PD-1 blockade"

**Supplemental Materials and Methods**

**Patient collection.** Patient samples were collected as part of a randomized clinical trial in which the patients (n = 3) participated in the nivolumab IV plus motolimod SC arm. Fresh tumor tissue biopsies were collected (Day 0) before two doses (Days 1 and 15) of intravenous (IV) anti-PD-1 blockade (nivolumab) and four doses (Days 1, 8, 15, and 22) of subcutaneous (SC) toll-like receptor 8 (TLR8) agonist (motolimod). Tumors were then resected on day 23 (Pt1_R), day 30 (Pt2_R), and day 28 (Pt3_NR). As this was considered secondary use of the patient samples, potentially identifying information such as patient weight and sex were not included as variables. Investigators were not blinded when using the patient samples.

**Multiplex Immunofluorescence (mIF) staining.** We used a commercially available staining kit (Opal 7-plex, Akoya, cat# NEL811001KT), and followed the manufacturer’s instructions. Nuclei were labeled using 4′,6-diamidino-2-phenylindole (DAPI, Spectral DAPI FP 1490, Perkin Elmer, Boston, MA, 1 ug/mL). Subsequently, a coverslip with mounting media (ProLong Glass Antifade Mountant, P36980 inner, refractive index 1.52, Invitrogen) was applied.

**Immunohistochemistry (IHC) validation.** To validate our multiplexed immunofluorescence staining, single sections of tumor were stained with chromogen-based IHC. All staining was manually performed, with antibodies against the following: pan-cytokeratin AE1/AE3 (Leica/(AE3/AE3)/Mouse-IgG1,  225 mg/L) CD19 (Leica/BT51E/Mouse-IgG2B, 35 mg/L), CD56 (Leica/CD564/Mouse, 11 mg/L), CD16 (Cell Signaling/D1N9L/Rabbit IgG, 100 µg/mL), alpha Smooth Muscle Actin (SMA) (Dako/1A4/Mouse-IgG2a, kappa 71 mg/L), Ki67 (Thermo Fisher/SP6/Rabbit-IgG,  0.029 mg/ml), CD8 (Cell Signaling/D1N9L/Rabbit IgG, 28.5 mg/L), CD4 (Abcam/EPR6855/ Rabbit monoclonal-IgG, 100 µg/mL), PD-1 (Abcam/EPR4877/ Rabbit monoclonal-IgG, 1.85 mg/ml), CD20 (Leica/L26/Mouse-IgG2A, kappa, 95 mg/L), LAMP3 (Thermo Fisher/ (PA5-84069) Polyclonal-Rabbit-IgG, 0.1 mg/mL), CD163 (Leica/10D6/Mouse, 49 mg/L), and TLR-8 (Abcam/44C143/Mouse monoclonal IgG1, 1 mg/mL). The specificity of each antibody was visualized using secondary labeling with a DAB (3,3'-diaminobenzidine) substrate kit (Vector Labs, SK-4100). This methodology uses a diaminobenzidine reaction to detect antibody labeling, and a hematoxylin counterstaining. A board-certified head and neck pathologist (Dr. Peter Sadow) reviewed and verified positive controls for each marker.

**Multispectral Imaging Platform.** All slides, including IHC, were imaged at 20X (NA 0.6, Vectra3, Perkin Elmer, bulb intensity 10%). This is an automated pathology imaging system that allows for the extracting of proteomic and morphometric information from intact FFPE tissue sections (chrome-extension://efaidnbmnnnibpcajpcglclefindmkaj/https://resources.perkinelmer.com/lab-solutions/resources/docs/prd_vectra3_012509_01.pdf). Together with Phenochart (version 1.1.0) and Inform 2.5.1, offers a user-friendly workflow for unique channel separation and analysis of up to seven markers in the same tissue section.

**Spectral Library Creation.** A spectral library was created according to Akoya Opal Assay Development Guide (https://www.akoyabio.com/wp-content/uploads/2020/04/Opal-Reagents_Brochure_Opal-Assay-Development-Guide.pdf ). Using HNSCC tissue, single-plex slides were created for each marker with a matched fluorophore, 4',6-diamino-2-phenylindole (DAPI), and one unstained slide for autofluorescence detection. All antibodies were tested for sensitivity and specificity to determine the optimal concentrations for primary antibodies. Further optimization was done until spectral readouts for each marker were between 5-30 normalized counts (InForm). In brief, representative areas from single-plex slides were captured at 20x magnification using the Vectra 3, followed by spectra extraction and storing using InForm 2.5.1. The quality of the spectral library was assessed using the R package vignettes/unmixing_quality_report.Rmd to measure unmixing quality. Raw images were spectrally unmixed using the generated libraries for later import into our data analysis platform (HALO Indica Labs).

**Isolation of tumor and immune cell infiltrates for scRNAseq.** Tissue was minced into small fragments inside a 1.5mL Eppendorf tube containing 420µL DMEM, 42µL of enzyme H, 21µL enzyme R, and 5µL of enzyme A (provided with the kit Miltenyi Biotec; Cat#130-095-929), followed by the addition of 512µL of DMEM, reaching a total volume of 1mL. The tube containing the small tissue fragments was briefly vortexed and placed in an Eppendorf thermomixer for 15 minutes at 400rpm, 37°C. At the end of the incubation, the dissociated tissue was placed over a 50µm filter (Sysmex; Cat# 04-004-2327), washed with 2mL of 4°C complete DMEM (containing 10% heat-inactivated FCS), minced with a 1mL syringe plunger, and washed with additional 5mL of complete DMEM. Dissociated cells were then filtered through a 30µm filter (Sysmex; Cat# 04-004-2326), and the tube was spun at 1500rpm at 4°C for 5 minutes. Next, the supernatant was removed, and ACK (Ammonium-Chloride-Potassium) cell lysing buffer (Gibco, cat#A1049201) was added to remove any red blood cells present in the pellet, and the tube was immediately spun at 1500 rpm at 4°C for 5 minutes, followed by removal of the ACK buffer. The sample was then resuspended in complete DMEM at 4°C, and the number of viable cells was determined using a hemocytometer (Bright-line; Cat# 1492), with cells labeled using trypan blue (Gibco, 0.4% solution, Cat#15250061). In all cases, the viability of dissociated single-cell suspensions was increased to >=90% by using a modified protocol of the EasySep Dead Cell Removal (Annexin V) kit (STEMCELL; cat#17899). First, cells were washed with 1mL of isolation media (PBSx1 + 2% FCS cat#16140-071 + 1mM $CaCl_2$), resuspended with 100µl of the same buffer, and placed into a PCR 8-strip well (cat#1402-4700) (1x10^6/well). Second, 10µl of dead cell removal (Annexin V) and biotin selection cocktails (both provided with the kit) were added, gently mixed, and incubated for 4 minutes at room temperature. Third, 20µl of vortexed RapidSpheres and 60µl of isolation media were added (total volume/well of 200 µl), mixed, and incubated for 3 minutes at room temperature. Fourth, the sample was placed on a 10x Genomics magnetic separator high-side (cat#120250) for 5 minutes. At the end of the incubation, the clear supernatant containing the live cells was moved into a new 1.5mL Eppendorf tube, spun at 1500rpm, 4°C, for 5 minutes, resuspended in complete DMEM and cell counts (concentration/mL) and viability was estimated using trypan-blue and a hemocytometer.

**Library preparation and 10X genomics sequencing**. Gene expression (GEX) and TCR libraries were generated using the 10x Genomics Chromium NextGEM Single Cell V(D)J Reagents kits v1.1 (PN-1000165; PN-1000020; PN-1000005; PN-1000120) according to the manufacturer instructions. We performed quality control evaluation for all samples after GEX and TCR cDNA and library generation, measuring the concentration (Qubit dsDNA High Sensitivity kit; Invitrogen; Cat#Q32854) and size distribution (Agilent High Sensitivity BioA

DNA kit; Cat#5067-4626). Samples were sequenced in pairs (pre/post) using a NextSeq 500 sequencer (Illumina).

**Processing of GEX and TCR scRNAseq.** Raw sequencing reads were aligned to the human reference genome hg38 (GRCh38-20202-A from 10X Genomics) and TCR VDJ reference (GRCh38-alts-ensembl-5.0.0 from 10X Genomics) using Cell Ranger (v6.1.1). Each sample's raw output, fastqs, was processed using Cell Ranger multi command with the –flags to expect 10,000 cells and include introns. Samples were then aggregated based on donors using Cell Ranger without normalization. Count matrices were loaded into Seurat (v3), and cells with fewer than 150 genes, or 15% of UMIs mapped to mitochondrial genes, were filtered. Potential doublets were filtered using DoubletFinder (chris-mcginnis-ucsf/DoubletFinder) with optimal pK values found through the mean-variance normalized bimodality coefficient.

**Clustering and annotation of tumor and tumor-infiltrating immune cells.** The resulting count matrix was further processed by normalizing the expression through Log normalization and scaled to center gene expression matrixes. The top 2000 highly variable genes based on the variance stabilizing method were determined and used to perform principal component analysis. To determine the dimensions used to perform clustering, Elbow plots and JackStraw analyses were performed. To perform clustering, the first 40 principal components were used to construct the nearest neighbor, and clusters were identified with the Louvain algorithm. To annotate the resulting 22 clusters, differential gene expression was performed with a log fold threshold of 0.25. Well-studied marker genes of immune cell types, exhaustion markers, and cytokines were used to assign predicted cell types. The uniform manifold approximation and projection (UMAP) was used to visualize the clusters.

**Sub-clustering and refinement of T cell and myeloid populations.** T cell and myeloid cell clusters gene expression and TCR clonotypes were extracted for further investigation. Separately, each cell type was normalized with a log normalization and scaled, and 2000 variable features were defined as described above. For T cells, the first 15 principal components were used to construct nearest neighbors and define clusters. For myeloid cells, the first 10 principal components were utilized. Clustering and annotation based on differential gene expression were repeated as described above.

**Copy number calling and phylogenetic tree construction.** Cells were classified based on CD45 expression with positive cells defined as having *PTPRC* counts greater than zero. InferCNV (InferCNV of the Trinity CTAT project: https://github.com/broadinstitute/InferCNV) was used to predict copy number alterations with CD45+ cell clusters as references. Copy number alterations were predicted using the Hidden Markov model (HMM) with a minimum average read count per gene set at 0.1 and the prediction of subclusters. Phylogenetic trees of tumor cells were then constructed using Uphyloplot2 (harbourlab/uphyloplot2) (Suppl. Figures 2-4).

**Processing, clustering and differential gene expression of Luoma et. al anti-PD-1 monotherapy treated HNSCC.** 10X genomics feature matrices from Luoma et.al [17], PD-1 monotherapy alone (n=3), were accessed from Gene Expression Omnibus (Accession GSE200996). Only patient data with samples collected before and after anti-PD-1 treatment were utilized. Cells that had fewer than 150 genes or 15% of UMIs mapped to mitochondrial genes were filtered using Seurat (v3). Potential doublets were filtered using DoubletFinder (chris-mcginnis-ucsf/DoubletFinder) with optimal pK values found through the mean-variance normalized bimodality coefficient. Filtered TCR alpha and beta contigs were then integrated by barcode using Seurat and scRepertoire (ncborcherding/scRepertoire) based on the VDJC genes and CDR3 nucleotide sequence. The resulting gene expression matrix and the original dataset described above were each normalized, and scaled, and 2000 variable features were defined as described above. The datasets were then integrated by ranking features by the number of datasets deemed variable and the top 2000 features. The integrated data set was scaled, and the first 40 principal components were used to identify clusters. To annotate clusters, differential gene expression analysis, as described above, was used in combination with integrating previously labeled clusters from Luoma et al. [17]. Only the samples with serial samples before and after anti-PD-1 treatment were utilized**:**

GSM6048143_raw_feature_bc_matrix_P23_pre-Tx,

GSM6048145_raw_feature_bc_matrix_P27_pre-Tx,

GSM6048146_raw_feature_bc_matrix_P29_pre-Tx,
GSM6048157_raw_feature_bc_matrix_P23_post-Tx,
GSM6048161_raw_feature_bc_matrix_P27_post-Tx,
GSM6048163_raw_feature_bc_matrix_P29_post-Tx

**Supplemental Figures:**

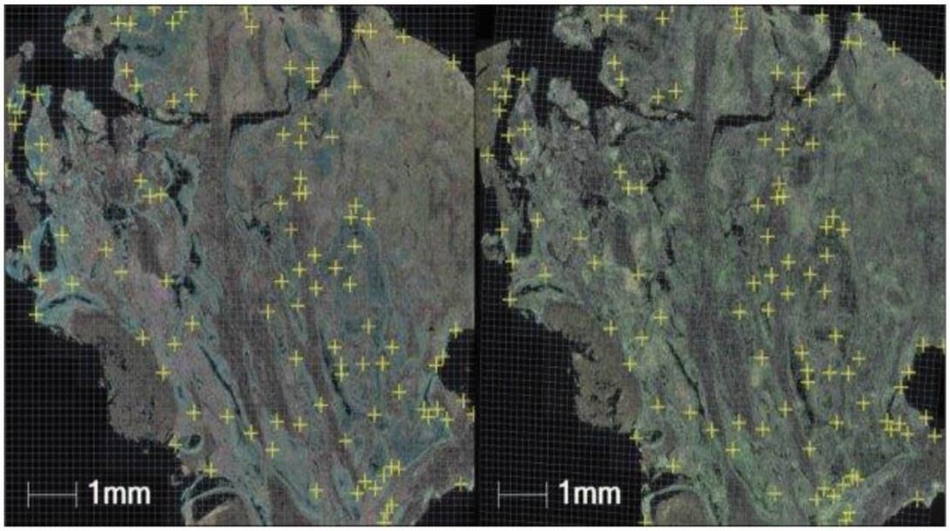


**Supplementary Figure 1: Registration marks for digital image fusion.** Two serially cut slides were fused after serial registration. Registration marks helped to improve the accuracy of serially fused images.

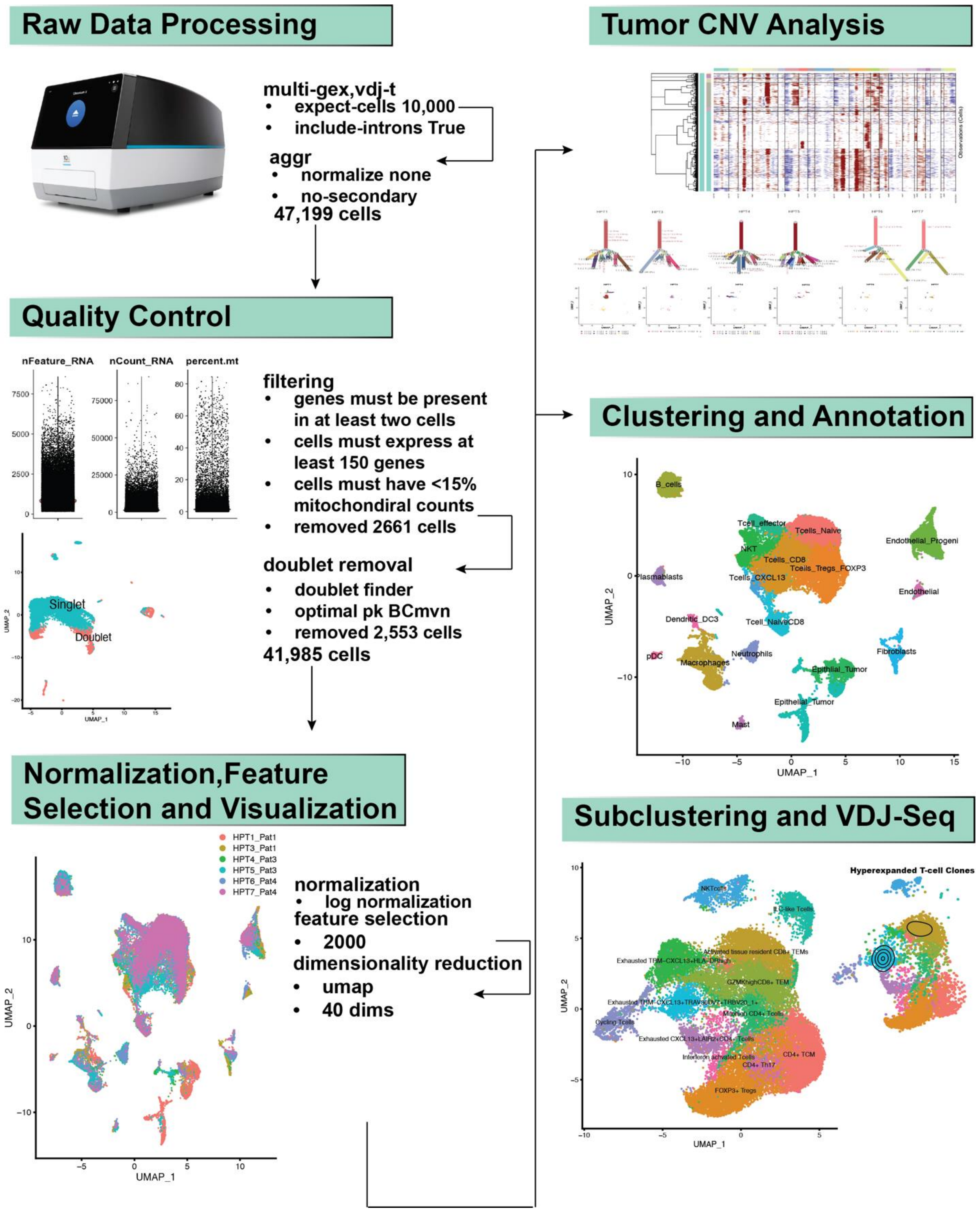


**Supplementary Figure 2. Overview of single-cell RNAseq processing workflow.** Single-cell RNAseq files were processed with cellranger with an expected cell count of 10,000 per sample.

Cells were quality-filtered based on mitochondrial-related gene counts and total gene count. Doubletfinder was used to remove potential doublets captured from droplet-based sorting. 40 dimensions and 2000 features were used for principal component analysis and UMAP clustering. InferCNV was used to determine potential copy number changes in tumor cells.

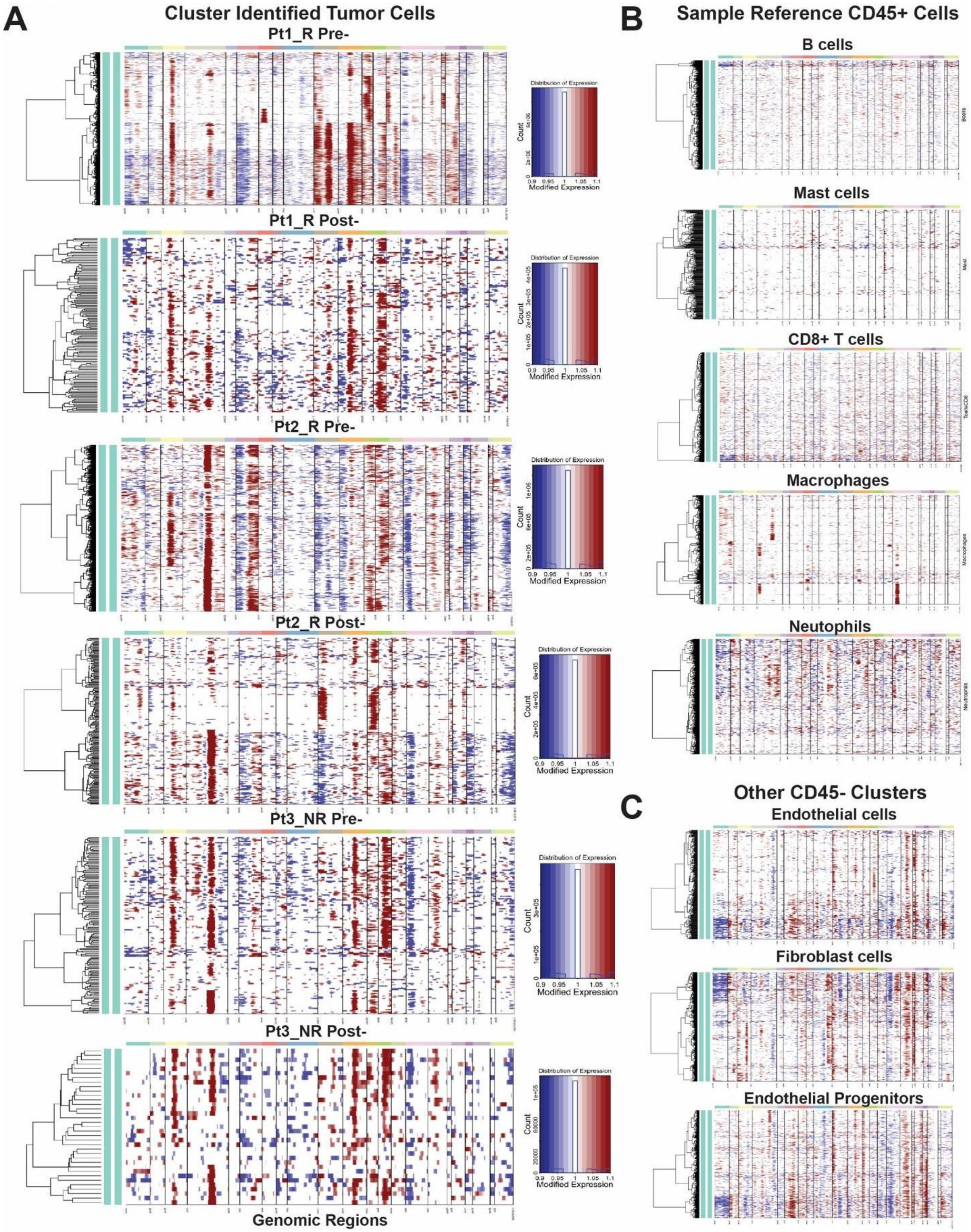
A
Cluster Identified Tumor Cells
Pt1_R Pre-
Pt1_R Post-
Pt2_R Pre-
Pt2_R Post-
Pt3_NR Pre-
Pt3_NR Post-
Genomic Regions
Distribution of Expression
Count
Modified Expression
B
Sample Reference CD45+ Cells
B cells
Mast cells
CD8+ T cells
Macrophages
Neutophils
C
Other CD45- Clusters
Endothelial cells
Fibroblast cells
Endothelial Progenitors

**Supplementary Figure 3. Copy Number Alterations inferred by InferCNV.** Cells were labeled as CD45+ or CD45- based on a minimum transcript count of one. Cells annotated as immune cell clusters and CD45+ were used as a reference to determine the copy number states of the remaining cells using inferCNV (https://github.com/broadinstitute/inferCNV) with the subclonal calculation option (cluster_by_groups=TRUE, denoise=TRUE, HMM=TRUE, analysis_mode=”subclusters”). **(A)** The expression values for each genomic region, representing a copy number state with gains in red and losses in blue, of cells previously annotated as epithelial_tumor cell clusters for each patient and sample are represented. Each row represents a cell, and each column is a genomic region sorted by chromosome and genomic location. **(B)** The resulting normalized expression values for each genomic region of 5 representative reference cell subsets. **(C)** The resulting normalized expression values for each genomic region of 3 non-immune cell subsets.

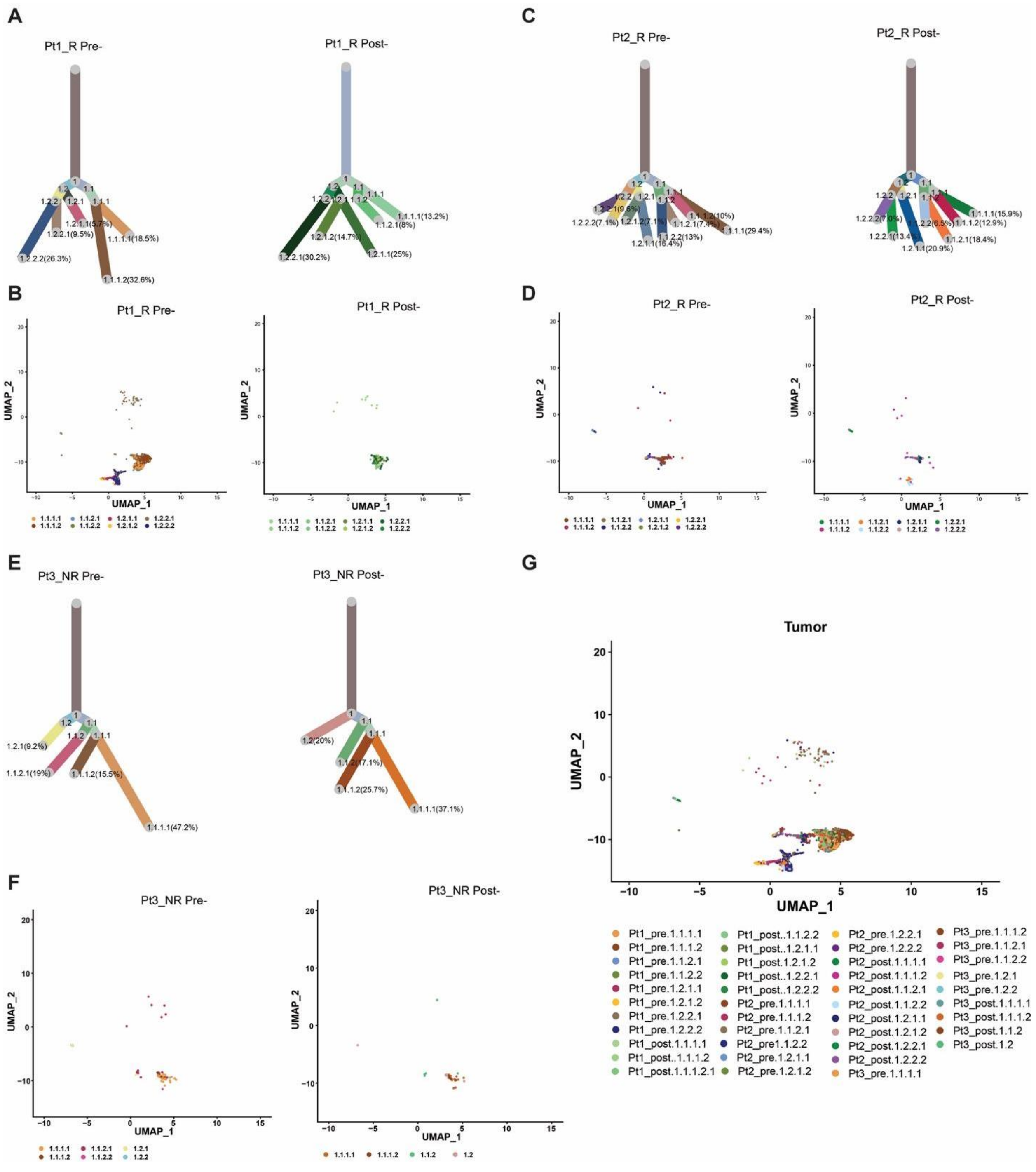


**Supplementary Figure 4. Tumor subclone populations and phylogenetic trees based on copy number profiles.** A phylogenetic tree was created using Uphyloplot2 (https://bmcgenomics.biomedcentral.com/articles/10.1186/s12864-021-07739-3) based on the percentage of cells within the subclusters identified by InferCNV. The length of each tree branch is proportional to the number of cells the clone has, with the original tree base representing a common progenitor cell. **(A)** Phylogenetic trees of patient 1 pre- and post-treatment. Each branch is labeled with the clone number determined by InferCNV and the percent of cells from that cluster. Clones under 5% of the population are not displayed. Through comparison of related copy number alterations across patient biopsy (Supplementary Figure 3), we determined branches in patient 1 post-treatment were phylogenetically related to patient 1 pre-treatment clone progenitor clones 1.1 and 1.1.1. **(B)** A UMAP of the epithelial tumor cells color-coded by subclone in the original UMAP space is displayed for patient 1 pre- and post-treatment biopsies. **(C)** Phylogenetic trees of patient 2 pre- and post-treatment with labeled subclones and percent of cells the subclone represented. **(D)** A UMAP of the epithelial tumor cells color-coded by subclone in the original UMAP space is displayed for patient 2 pre- and post-treatment biopsies. **(E)** Phylogenetic trees of patient 3 pre- and post-treatment with labeled subclones and percent of cells the subclone represented. Through comparison of related copy number alterations across patient biopsy (Supplementary Figure 3), we determined that three branches in patient 3 post-treatment were phylogenetically related to patient 3 pre-treatment clone progenitor clones 1.1 **(F)** A UMAP of the epithelial tumor cells color-coded by subclone in the original UMAP space is displayed for patient 3 pre- and post-treatment biopsies. **(G)** A UMAP of the epithelial tumor cells color-coded by subclone in the original UMAP space is displayed for all 6 patient biopsies.

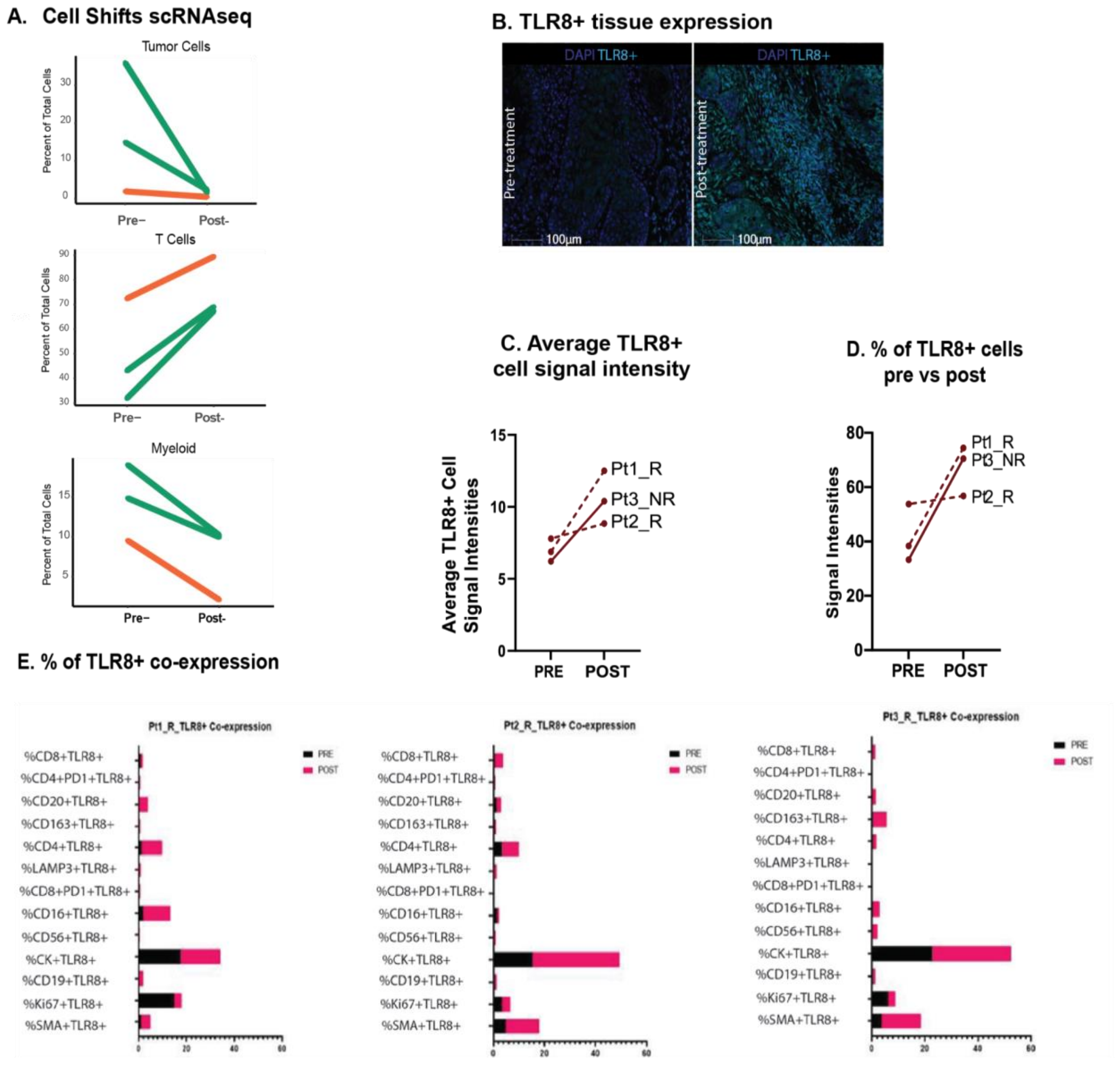


**Supplementary Figure 5**. **Cell shifts via scRNAseq and TLR8 protein expression changes after anti-PD-1 and TLR8 agonism. (A)** Single-cell RNA shifts from pre- to post-treatment in tumor, T cells, and myeloid cells. **(B)** TLR8 protein detection via mIF pre- and post-treatment. **(C)** Average signal intensities for TLR8+ cells. **(D)** Percentage of TLR8+ positive pre- and post-treatment. **(E)** Percentage of TLR8 co-expression with other markers in the panel.

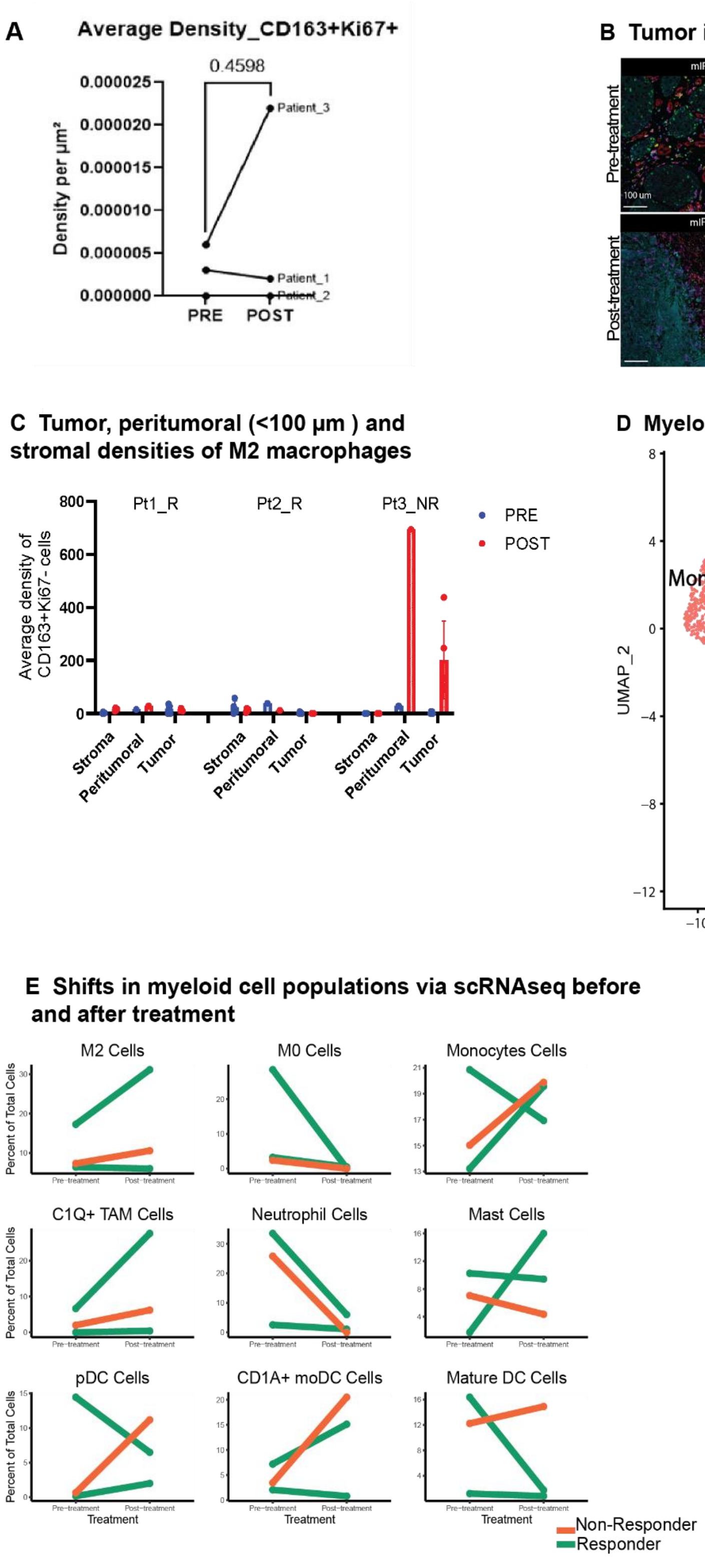
A
Average Density_CD163+Ki67+
0.4598
Density per µm²
Patient_3
Patient_1
Patient_2
PRE
POST
B Tumor infiltrating CD163+ cells
Pre-treatment
Post-treatment
mIF
Tumor Stroma
CD163 Cytokeratin Ki67 DAPI
100 um
C Tumor, peritumoral (<100 µm ) and stromal densities of M2 macrophages
Average density of CD163+Ki67- cells
Pt1_R
Pt2_R
Pt3_NR
PRE
POST
Stroma
Peritumoral
Tumor
D Myeloid cell populations via scRNAseq
UMAP_1
UMAP_2
Monocyte
M2
Neutrophil
C1Q+ TAM
M0
Tcell/mono
Tcell/neutro
CD1A+ moDC
Mast
Tcell/mast
Mature DC
pDC
Neutro
E Shifts in myeloid cell populations via scRNAseq before and after treatment
M2 Cells
M0 Cells
Monocytes Cells
C1Q+ TAM Cells
Neutrophil Cells
Mast Cells
pDC Cells
CD1A+ moDC Cells
Mature DC Cells
Percent of Total Cells
Pre-treatment
Post-treatment
Treatment
Non-Responder
Responder

**Supplementary Figure 6. Analysis of myeloid cells pre-and post-treatment. (A-C)** The density of M2 (CD163+) macrophages pre- vs post-treatment, with a marked increase in density in the non-responder. **(D)** UMAP shows clustering of all myeloid lineage cells. **(E)** Percentages of total cells for specific clusters per patient per time point.

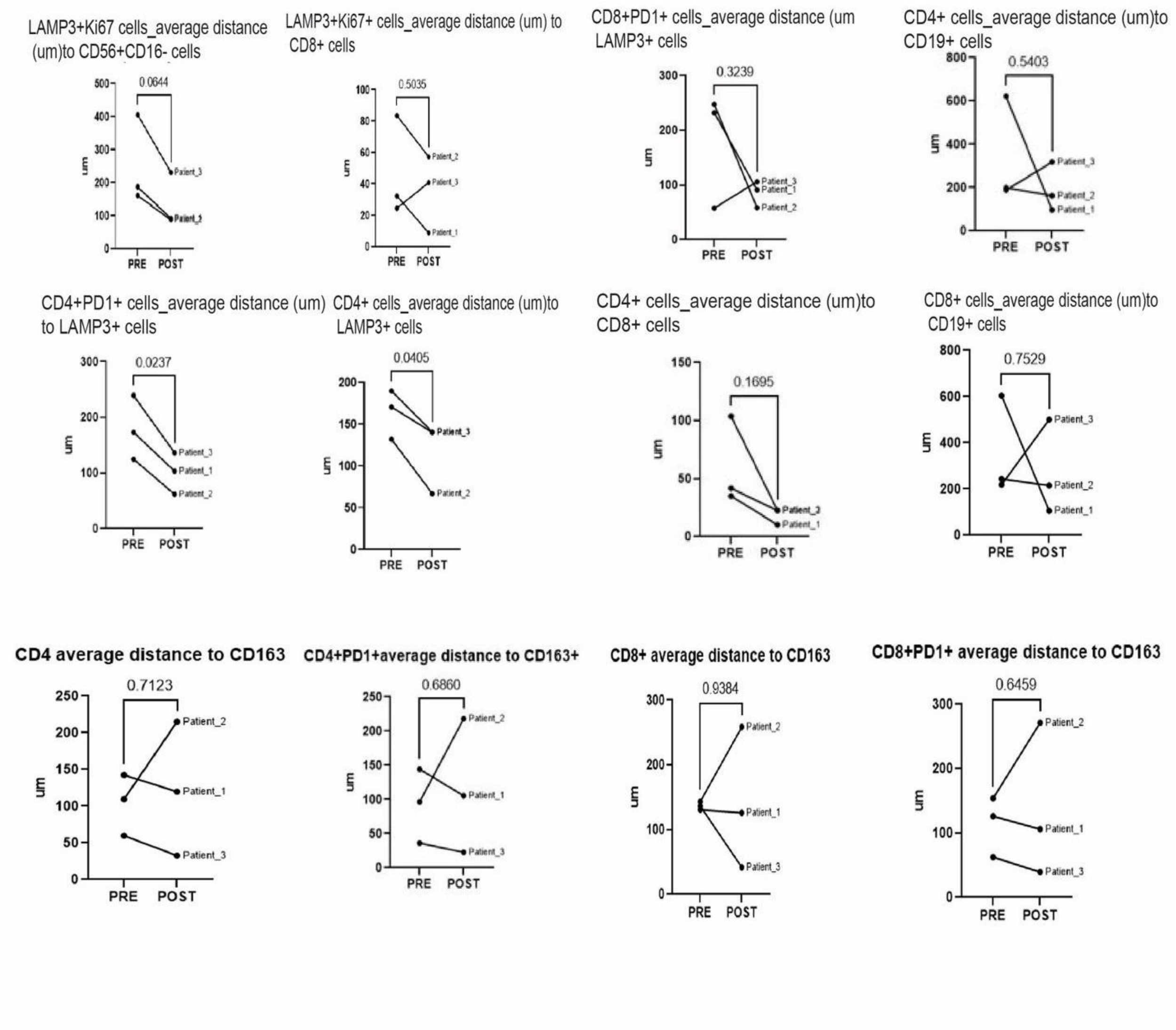


**Supplementary Figure 7. Proximity analysis.** Proximity analysis between proliferative dendritic cells (LAMP3+Ki67+) and NK cells (CD56+CD16-), T helper cells (CD4+), cytotoxic T cells (CD8+), exhausted T cells (PD-1+), B cells (CD19+), and M2 macrophages (CD163+).

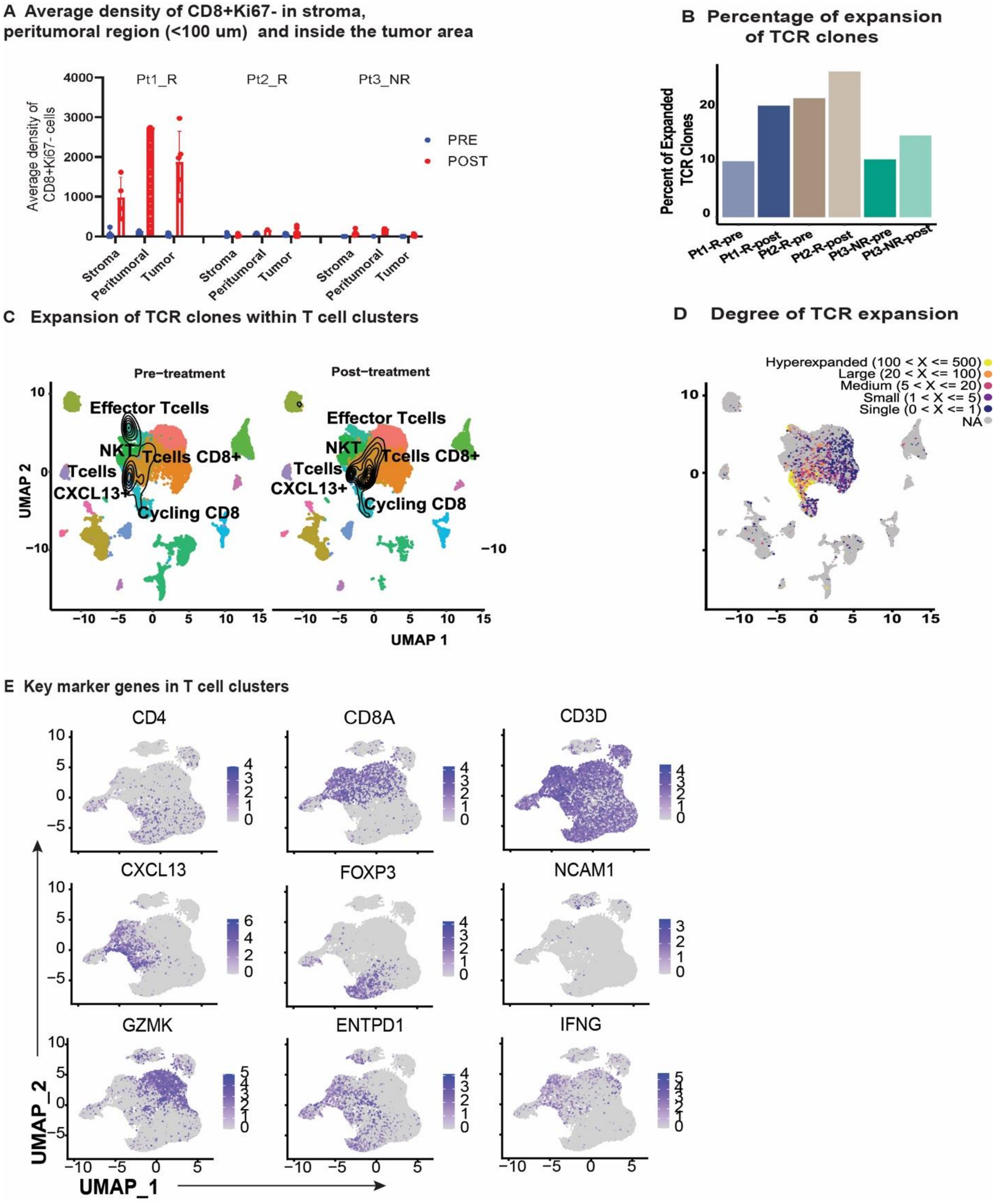
A Average density of CD8+Ki67- in stroma, peritumoral region (<100 um) and inside the tumor area
Pt1_R
Pt2_R
Pt3_NR
Average density of CD8+Ki67- cells
PRE
POST
Stroma
Peritumoral
Tumor
B Percentage of expansion of TCR clones
Percent of Expanded TCR Clones
Pt1-R-pre
Pt1-R-post
Pt2-R-pre
Pt2-R-post
Pt3-NR-pre
Pt3-NR-post
C Expansion of TCR clones within T cell clusters
Pre−treatment
Post−treatment
Effector Tcells
NKT
Tcells CD8+
Tcells CXCL13+
Cycling CD8
UMAP 2
UMAP 1
D Degree of TCR expansion
Hyperexpanded (100 < X <= 500)
Large (20 < X <= 100)
Medium (5 < X <= 20)
Small (1 < X <= 5)
Single (0 < X <= 1)
NA
E Key marker genes in T cell clusters
CD4
CD8A
CD3D
CXCL13
FOXP3
NCAM1
GZMK
ENTPD1
IFNG
UMAP_2
UMAP_1

**Supplementary Figure 8. T cell expansion after combinatorial therapy. (A)** The average density of cytotoxic T cells (CD8+Ki67-) in the stroma, peritumoral region (< 100 µm), and intratumoral region **(B)** Percentage of expansion of T-cell receptor clones. **(C)** UMAP shows the expansion of TCR clones within T clusters. **(D)** Density heatmap showing the degree of TCR expansion. **(E)** Key markers genes found in T cell clusters.

**A Random forest classifier trained with the pathologist annotations for TLS detection**

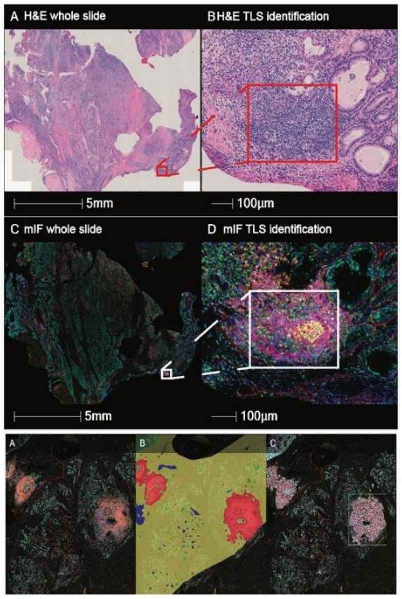


**D Percentage of contribution to TLS per marker**

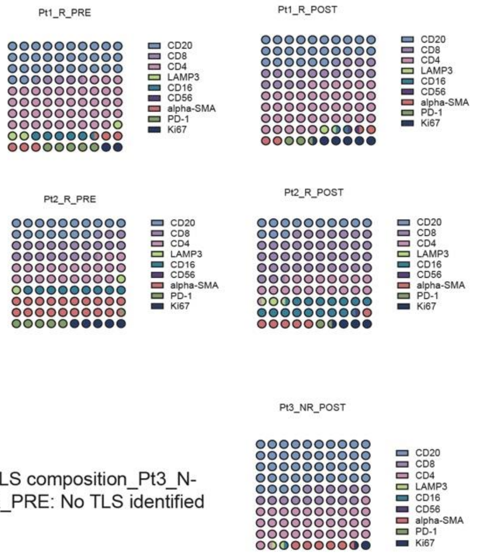


TLS composition_Pt3_N-R_PRE: No TLS identified

**B Identification of TLS (dots) in both pre and post treatment**

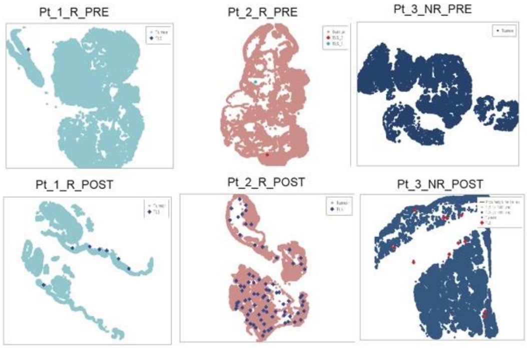


**E**

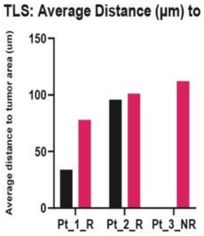


**C Number of TLS normalized by size, assesed via H&E by a pathologist**

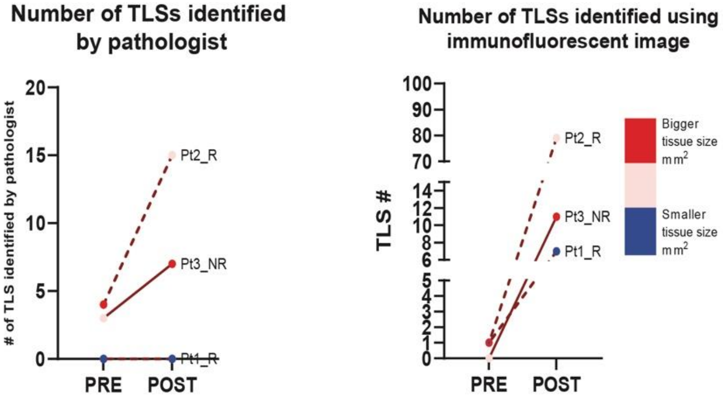

**Supplementary Figure 9. Analysis of Tertiary Lymphoid Structures. (A)** Identification of TLSs on serially cut H&Es by a pathologist (TLS/$mm^2$). **(B)** Graphical representation of TLS in the tissue section. **(C)** The raw TLS number was identified on H&E by the pathologist, and the raw number was identified via mIF. **(D)** Components per patient pre- and post-treatment. **(E)** Distance of TLS from tumor area per patient pre- and post-treatment.

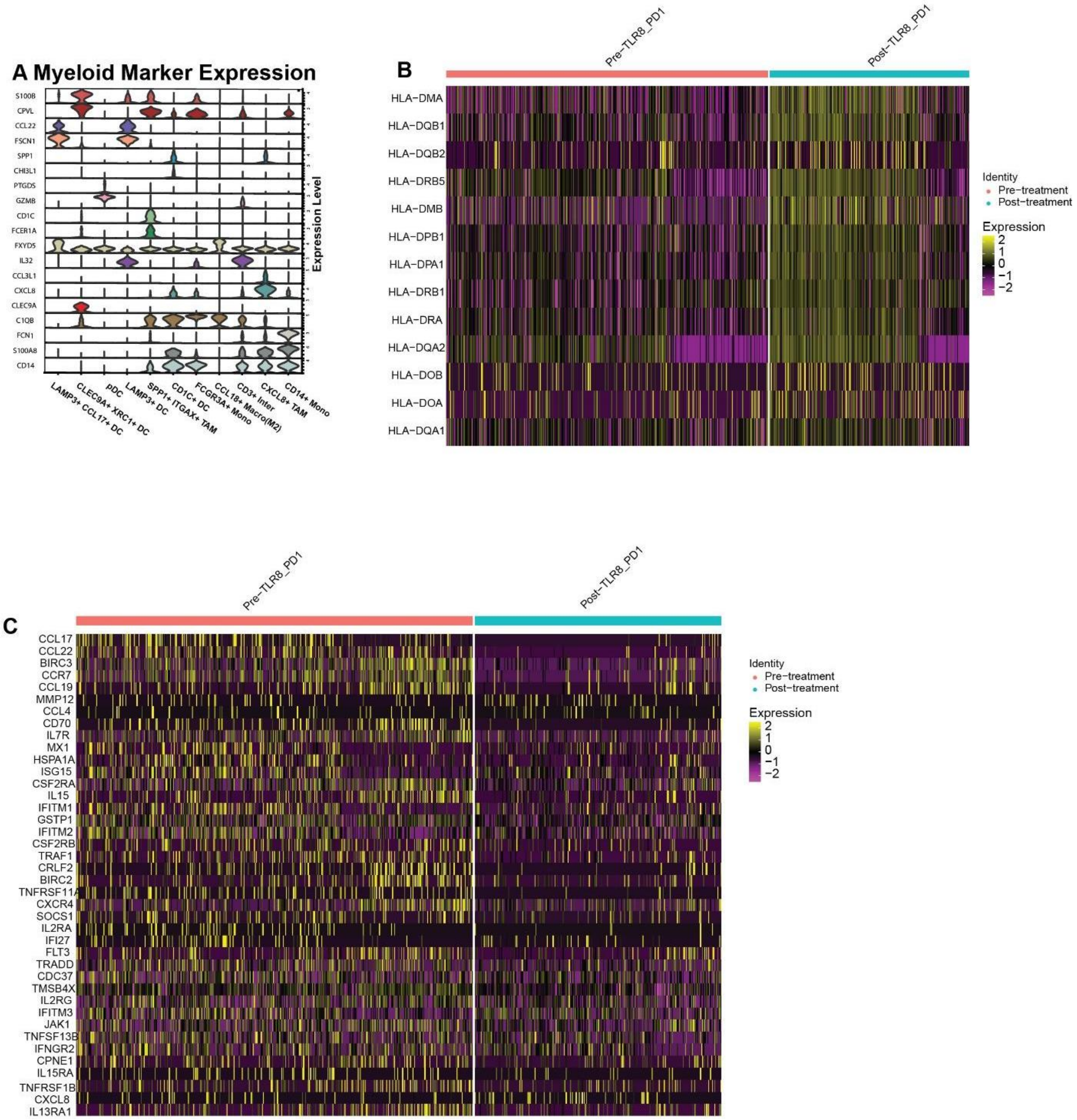


**Supplementary Figure 10. Dendritic Cell Changes with PD-1 blockade and TLR8 Agonism** **(A)** Myeloid marker expression. Two tissue-associated macrophage populations were defined by ITGAX expression, cytokine IL-8 producing gene CXCL8, and EMT-related gene SPP1. **(B and C)** Seurat was used to determining differentially expressed genes in non-plasmacytoid dendritic cells between pre-and post-treatment. Geneset enrichment was used, based on the log fold change, to associate these genes with the GO Biological Processes. The leading edge genes for GOBP_PEPTIDE_ANTIGEN_ASSEMBLY_WITH_MHC_CLASS_II_PROTEIN_COMPLEX and GOBP_CYTOKINE_MEDIATED_SIGNALING_PATHWAY are plotted in a scaled gene expression heatmap, respectively.
